# Noradrenalin induced the proliferation and migration of human pulmonary artery smooth muscle cells via endothelin-1

**DOI:** 10.64898/2026.08.25.747161

**Authors:** Ting-An Yen, Hsin-Chung Huang, En-Ting Wu, Heng-Wen Chou, Hung-Chieh Chou, Chien-Yi Chen, Shu-Chien Huang, Yih-Sharng Chen, Po-Nien Tsao, Fu-Shan Jaw, Ching-Chia Wang

**Author notes:** **Corresponding author:** Ching-Chia Wang, **Address:** No.8, Chung Shan S. Rd.(Zhongshan S. Rd.), Zhongzheng Dist., Taipei City 10041, Taiwan (R.O.C.) **Email address:**.

## Abstract

**Background:** Pulmonary arterial hypertension (PAH) is a serious disease with poor prognosis, especially in infants or preterm babies and there is still no optimal treatment for this disease. Noradrenalin (NE) is a vasoactive mediator which is released by sympathetic ganglion. According to previous studies, NE/α1-adrenoreceptors is not only in regulating normal physiologic responses, but also in the pathogenesis of PAH. However, the mechanisms of NE in PAH are not fully understood.

**Methods:** Human PASMC (PASMC) was used in this study. Cell viability assay and Wound healing assay were used to evaluate the proliferation and migration of PASMC. Immunoprecipitation and western blots analysis were used to investigate the mechanisms which involved in NE-induced PASMC proliferation.

**Results:** We investigated that NE could induce human PASMC proliferation and migration. Furthermore, we first find that endothelin 1 (ET-1) signaling pathway plays an important role in NE-induced PASMC proliferation. ET1 is a critical molecular which is known for regulating cell growth and migration. We investigated that NE could increase NE-1 secretion, further enhancing ET-1 bind to its receptors. For further clarifying the downstream signals in NE/ET-1 induced PASMC proliferation, we detected the phosphorylation and expression levels of ERK and JNK.

**Conclusions:** By combining the results from ours and previous studies, we believed that JNK/c-jun pathway may play an important role in NE-induced PASMC proliferation.

## Introduction

Pulmonary arterial hypertension (PAH) is a serious disease with poor prognosis, especially in infants or preterm babies. There are lots of new treatment strategies developed in the past ten years. However, there is still no optimal treatment for this disease. In children, the disease burden is increasing due to the improved survival rate of children with congenital heart disease and premature infants. For premature infants, hypoxia with chronic lung disease is a major risk factor leading to pulmonary hypertension. In the development of PAH, oxygen (O_2_) plays an essential role in determining pulmonary vascular tone. Increasing O_2_ tension with the onset of air-breathing, pulmonary blood flow increases and pulmonary artery pressure falls compared with systemic levels over the first 24 hours of postnatal life^1^. It means that pulmonary vascular tone remains low even as the pulmonary circulation responds to compromised ventilation with vasoconstriction to prevent intrapulmonary shunting^2^. Therefore, the pulmonary circulation remodels which lead to right heart failure and even death.

According to previous studies, there are complex mechanisms involved in infant PAH development. Multiple lines of evidence point to a central role for the O_2_-sensitive transcription factor, hypoxia-inducible factor-1α (HIF-1α), in the regulation of pulmonary vascular tone^3–6^. According to an animal model with smooth muscle cell (SMC)-specific deletion of HIF-1α, HIF-1α plays a role in maintaining low pulmonary vascular resistance^7^. In another report, conditional SMC-specific deletion of HIF-1α mitigated hypoxic pulmonary hypertension. ET-1 is another important participant in PAH development^4^. Several studies indicated ET-1 could enhance pulmonary artery smooth muscle cells (PASMC) proliferation or inhibit PASMC apoptosis, which are critical processes in vascular remodeling during PAH development^8–10^. Some evidence showed that blocking the signal transduction of ET-1 could attenuate the vascular remodeling in PAH, indicating that ET-1 plays an important role in the PAH development^11^. The correlation between HIF-1α and ET-1 in PAH development was also discussed. ET-1 is thought to have the ability to induce HIF-1α expression in PASMC^12, 13^. Furthermore, in the previous study, we found that loss of HIF-1α increases microRNA (miRNA)-543 which decreases Twist expression, leading to an increase in ET-1 expression in PASMC. However, even though the molecular mechanisms for PAH development are more and more clear, there are still other important pathways that could control these critical signals for PAMSC proliferation and trigger PAH.

The effects of vasoactive mediators have also been known to play an important part in the progression of PAH. The previous study demonstrates that direct sympathetic ganglion blockage (SGB) achieved by injecting local anesthetic into the superior cervical ganglion attenuated monocrotaline-induced progression of PAH in rats^14^. The reason is due to the inhibition of arginase and oxidative stress and elevation of eNOS. Direct SGB could be applied as a novel therapeutic treatment modality for many cardiovascular diseases associated with endothelial dysfunction including PAH^14^. As we know, noradrenalin (NE) is the vasoactive mediator which is released by sympathetic ganglion. Therefore, it indicates the importance of NE and α1-adrenoreceptors in the pulmonary vasculature, which is not only in regulating normal physiologic responses, but also in the pathogenesis of PAH. α1-Adrenoreceptors in the pulmonary arteries have greatly increased affinity and responsiveness with their agonists when compared to other vessels^15^. The downstream signaling events in α1-adrenergic stimulation are an increase in calcium levels and the activation of protein kinase C, which mediate vascular contractile and proliferative responses^16^. The excessive stimulation of α1-adrenergic receptors produces smooth muscle contraction, proliferation, and growth.

Some data have proved the role of NE for vascular smooth muscle cells proliferation during PAH development. One study showed that NE could induce the proliferation of rabbit vascular smooth cells through small extracellular vesicles released by fibroblasts^17^. It is indicated that the activation ofα1-adrenergic receptors-related pathway by extracellular NE may be not the only mechanism for controlling NE-induced smooth muscle cells proliferation. The crosstalk between different cell types also play an important role in the effects of NE. Furthermore, P38 pathway was considered to participate in NE-induced rat PASMC proliferation^18^. Although these mechanisms could help explain the part of action mode of NE in promoting PASMC proliferation, there still have some critical mediators needed to be clarified for comprehensive understanding the role of NE in Human PASMC proliferation. From our perspective, it is important to know which secreted molecules are responsible for communicating with PASMCs and which receptor is activated and responsible for executing PASMC proliferation. The information could help us utilize the NE pathway to establish more comprehensive treatment strategies for PAH.

Endothelin 1 (ET1), 21-amino acid peptide secreted from endothelium, epithelium and vascular smooth muscle cells, is a critical molecular which could regulate cell growth and migration, including the PASMC proliferation during PAH development^8, 19–22^. Our previous study also indicated miRNA-486-5p could regulate PASMC proliferation by ET-1^23^. ET1, secreted by endothelium cells or other cell types, could bind the G-protein -coupled receptors (ETA and ETB) and mediates the vasoconstriction and tissue repair^24^. Weng et al. demonstrated the ET1 induced human lung fibroblasts differentiation through ETAR/JNK/AP-1 signaling pathway^25^. ET1 is known to play a pivotal role in the pathobiology of PAH. ET1 also activated the mitogen-activated protein kinases (MAPK) and nuclear factor c-Jun in canine pulmonary smooth muscle cells^26^. The antagonists of ET1 receptors are widely used clinically but the therapeutic efficacy has been limited^27^. To sum up, it is known that both NE and ET-1 could enhance PASMC growth. However, whether NE is participated in the ET1-regulated PASMC proliferation is still unclear. In this study, we try to further clarify the mechanisms of NE on ET-1 related PASMCs proliferation and detect possible downstream signals in this process.

## Methods

### Cell culture

Human PASMC (PASMC) was purchased from Lonza (Basel, Switzerland). Cells were cultured in medium provided by Lonza added with 5% FBS, 0.2% rhFGF-B, 0.1% rhEGF, 0.1% Gentamicin sulfate, Amphotericin-B, and 0.1% Insulin. Then cells were put into a 37℃ humidified incubator with 5% CO_2_. Only the Cells from passages 4–8 were used for the experiments in this study.

### ET-1 ELISA

Secreted ET-1 was measured using an ET-1 colorimetric immunometric ELISA kit purchased from Enzo Life Sciences (NY, USA). After PASMC was treated with 0.1 and 1 μM NE for 6hr, the growth media were collected. At the same time, PASMC was lysed by RIPA and total cell lysates were collected for total protein concentration measurement. Secreted ET-1 in growth medium was measured according to the manufacturer’s protocol. Optical density was measured at 450 nm and the concentration of ET-1 in samples were calculated using a standard curve established by different concentrations of recombinant ET-1. Final data were normalized to total protein concentration.

### Cell viability assay

PASMC were incubated into 96-well and treated with 0.1-10 μM NE for 6hr. After NE treatment. PASMC were rinsed by 200 μL PBS two times to remove the residual growth medium containing NE. 200 μl serum-free medium with 3-(4,5-dimethyl thiazol-2-yl)-2,5-diphenyltetrazolium bromide (MTT) (MilliporeSigma, MA, USA) was add into each well, and PASMC was incubated under 37℃ with 5% CO_2_ for 2hr. After incubation, PASMC were rinsed by 200 μl PBS two times again to remove the residual medium containing MTT. 200 μl dimethyl sulfoxide was added in to each well and the whole 96-well culture plate was gently shaken for 15 minutes. The optical density of 570 nm was measured to calculate cell viability.

### Wound healing assay

PASMC were incubated into 24-well until a monolayer formed. To create the wound gap, the sterile pipette tip was used. The monolayer cells were scratched using the sterile pipette tip from the same direction. After scratching, cells gently were rinsed two times with PBS to remove the detached cells, followed by the growth medium replacement. The scratched PASMC were treated with 1 μM NE and incubated for 6hr. The cell migration rate was evaluated by calculating the mean distance of cells migrated into the wound area.

### Western blot analysis

After 0, 0.1 or 1μM NE treatment for 6hr, PASMC were homogenized in 150 μl RIPA cell lysis buffer containing proteinase inhibitors. Then, the cell lysates were centrifuged at 13,000 g for 30 min at 4℃ to collect protein suspensions. The 30–80μg total protein were separated by electrophoresis using 8 or 10% sodium dodecyl sulfate polyacrylamide gel, followed by transferred the protein to polyvinylidene difluoride membranes. After blocked with phosphate-buffered saline with Tween 20 (PBST) buffer with 5% skimmed milk powder, the membranes were immersed in the PBST buffer containing antibodies for ET-1, c-Jun, p-JNK, JNK, ERK, p-ERK, androgen receptor (ARα), endothelin-1 receptor A (ETRA), endothelin-1 receptor B (ETRB), or β-actin (Cell Signaling Technology, MA, USA) overnight. After the membranes were washed by PBST two times, they were immersed in PBST buffer containing anti-mouse or anti-rabbit secondary antibodies for 1 hr. After washed by PBST two times again, the chemiluminescence of antibody-reactive bands were enhanced by ECL kits (Merek Millipore, Hesse, Germany) and detected by the X-ray film (Kodak, NY, USA). Densitometric analysis was performed by Image-J.

### Immunoprecipitation

After treatment, equal amount of proteins from cell lysate were bound using an immunoprecipitation antibody (ETBR or ETAR) over night, and immunoprecipitated with adding protein A beads (Blossom Biosciences LTD.) for 2 hours at 4°C. The immunoprecipitate was collected by centrifuge at 12,000 x g for 20 seconds to discard the supernatant and then washed three times with cold lysis buffer and subjected to immunoblot analysis.

### Statistics

Results are expressed as means ± SEM according to the experiments repeated with a minimum of three times. Statistical significance was assessed with Student’s t-test. A p value of < 0.05 was considered as the statistical significance.

## Results

It is known that pulmonary artery smooth muscle cells (PASMC) proliferation is a key pathological characteristic during PAH development. According to previous studies, we know that NE could enhance vascular smooth muscle cells in rats and rabbits. However, whether NE has the same effects on human PASMC is still unclear. In this study, we wanted to confirm the effects of NE on primary human PASMC first. PASMC were treated with 0, 0.1, 0.5, 1, 5, 10 μM NE for 24hr, and the cell viability were evaluated by MTT assay. From figure1, we found that NE could increase cell viability significantly in all NE concentrations. Furthermore, the effect of NE on increasing cell viability is strongest in 0.5 and 1 μM NE group, but weaker in 5 and 10 μM NE group. According to the results, we used 0.1 or 1 μM NE for the following experiments. We noticed that NE also could enhance the migration of PASMC. In figure 1B and 1C, 1 μM NE promoted PASMC migration significantly in 6hr detecting by wound healing assay.

**Figure 1.**
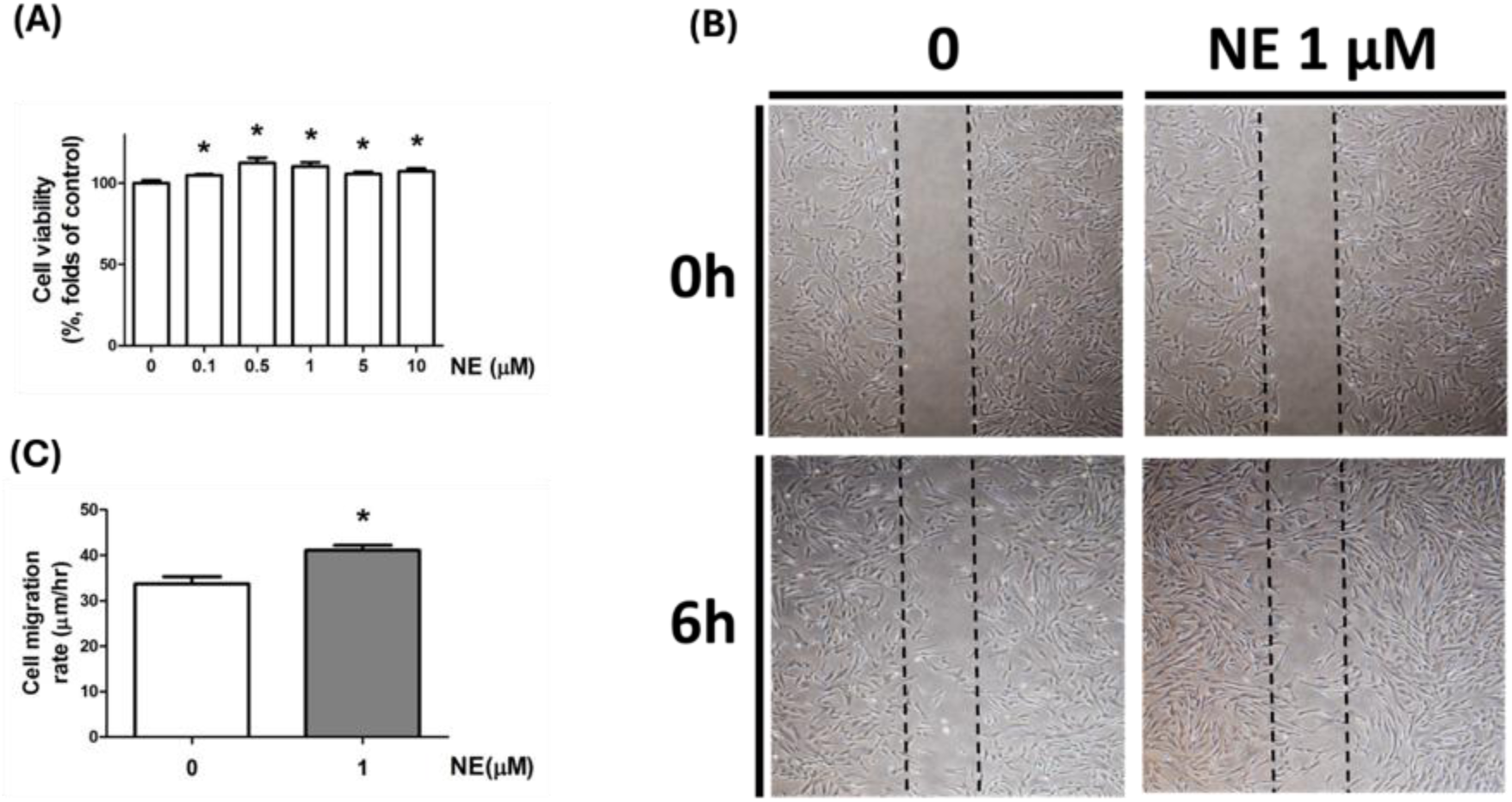
Noradrenalin (NE) induced PASMC proliferation and migration. After 0.1-10 μM NE treatment for 6hr, the cell viability of PASMC was evaluated by MTT assay (A). The migration ability of PASMC which was induced by 1 μM was evaluated by wound healing assay (B). The cell migration rate was evaluated by calculating the mean distance of cells migrating into the wound area (C). Results are expressed as means±SEM for at least three independent experiments. *P < 0.05 compared to control group.

ET-1 is a known factor which regulates the PASMC proliferation during PAH progression. We want to figure out the effects of NE on ET-1 expression and secretion, thus we treated PASMC with 0.1-10 μM NE for 6hr and detected ET-1 expression by western blots. In figure 2, we observed that ET-1 expression increased strongly on 0.5 and 1 μM NE treatment group. However, we cannot find ET-1 upregulation effect on 5 and 10 μM NE treatment group. The result was consistent with the result of cell viability. Combining the results from figure 1 and 2, we hypothesized NE may regulate multiple signal pathways to enhance PASMC proliferation. ET-1 pathway plays an important role for enhancing cell viability induced by lower concentrations of NE (0.1-1 μM NE). However, treated with higher concentrations of NE (5 and 10 μM NE) could not induce ET-1 expression. On the same concentrations of NE (5 and 10 μM NE), the increase of cell viability was weaker compared with 0.1-1 μM NE treatment group. Treated with 1 μM NE also could promote ET-1 secretion significantly. This result showed that NE may cause a paracrine effect for enhancing PSAMC proliferation through ET-1 secretion.

**Figure 2.**
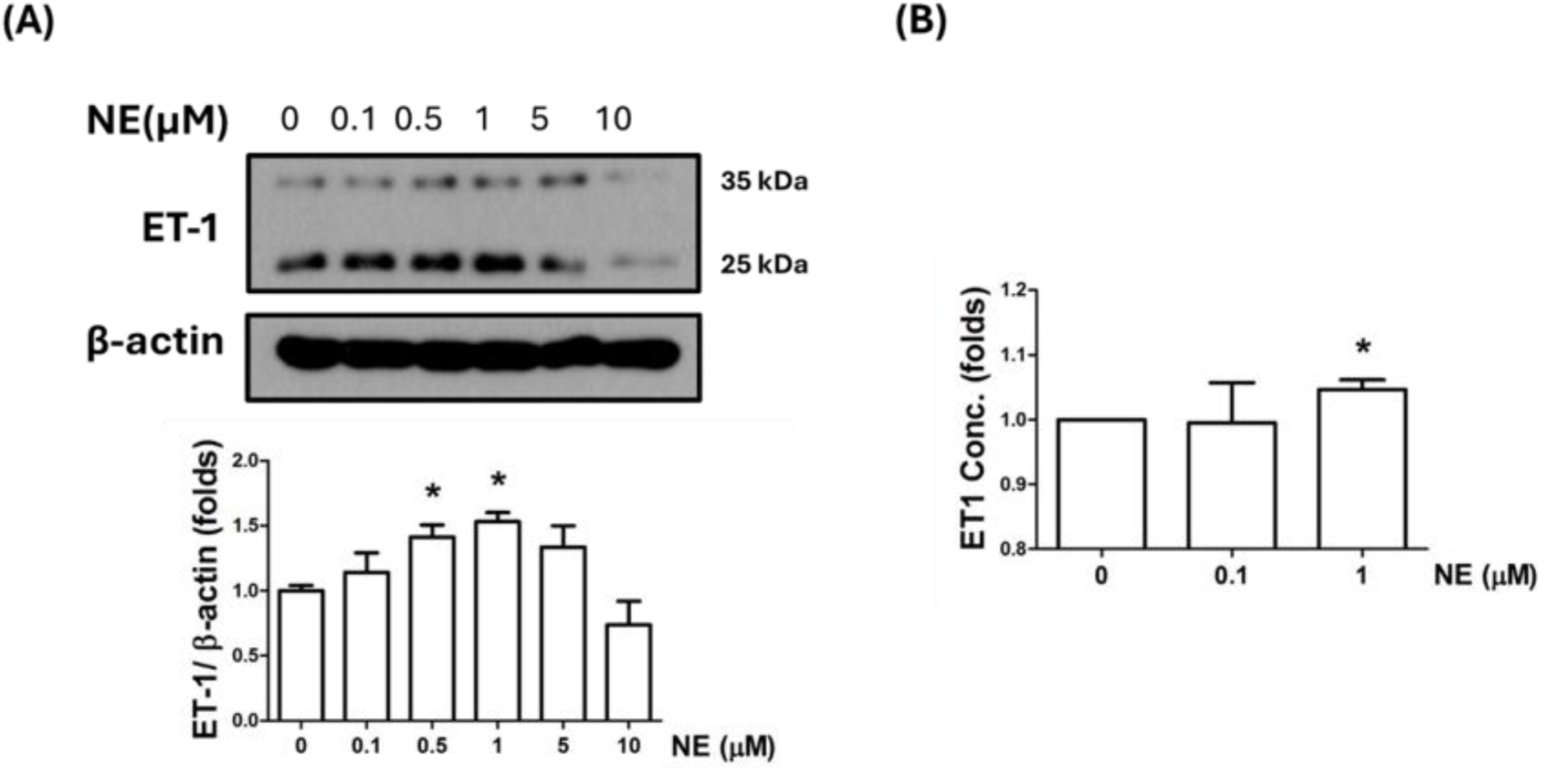
Noradrenalin (NE) increased the expression and secretion of ET-1. After 0.1-10 μM NE treatment for 6hr, the protein expression of ET-1 was evaluated by western blots (A). The secreted ET-1 in culture medium which was induced by 0.1 and 1 μM was measured by ELISA (B). Results are expressed as means±SEM for at least three independent experiments. *P < 0.05 compared to control group.

For further demonstrating the paracrine effect caused by NE, we try to figure out whether Endothelin A receptor and Endothelin B receptor (ETAR and ETBR), the two major kinds of receptor for secreted ET-1, were activated after NE treatment. After 0.1 and 1 μM NE treatment, the levels of androgen receptors alpha (ARα), one major kind of NE receptor, ETAR and ETBR were detected by western blots. We found that the levels of ARα and ETAR decreased in a dose dependent manner and exist a significant effect under 1 μM NE treatment (Figure 3). However, the level of ETBR was not affected by NE treatment (Figure 3). For understanding the interactions between ET-1 and its receptors (ETAR and ETBR), we conducted the immunoprecipitation by ETAR and ETBR antibodies and detected the level of ET-1 by western blots. In figure 4, we found that treatment with 1 μM NE caused an increase in the amount of ET-1 that binds to both ETAR and ETBR. These results showed that the NE-induced ET-1 secretion could activate both ETAR and ETBR pathway, which could further enhance PASMC proliferation by a paracrine effect.

**Figure 3.**
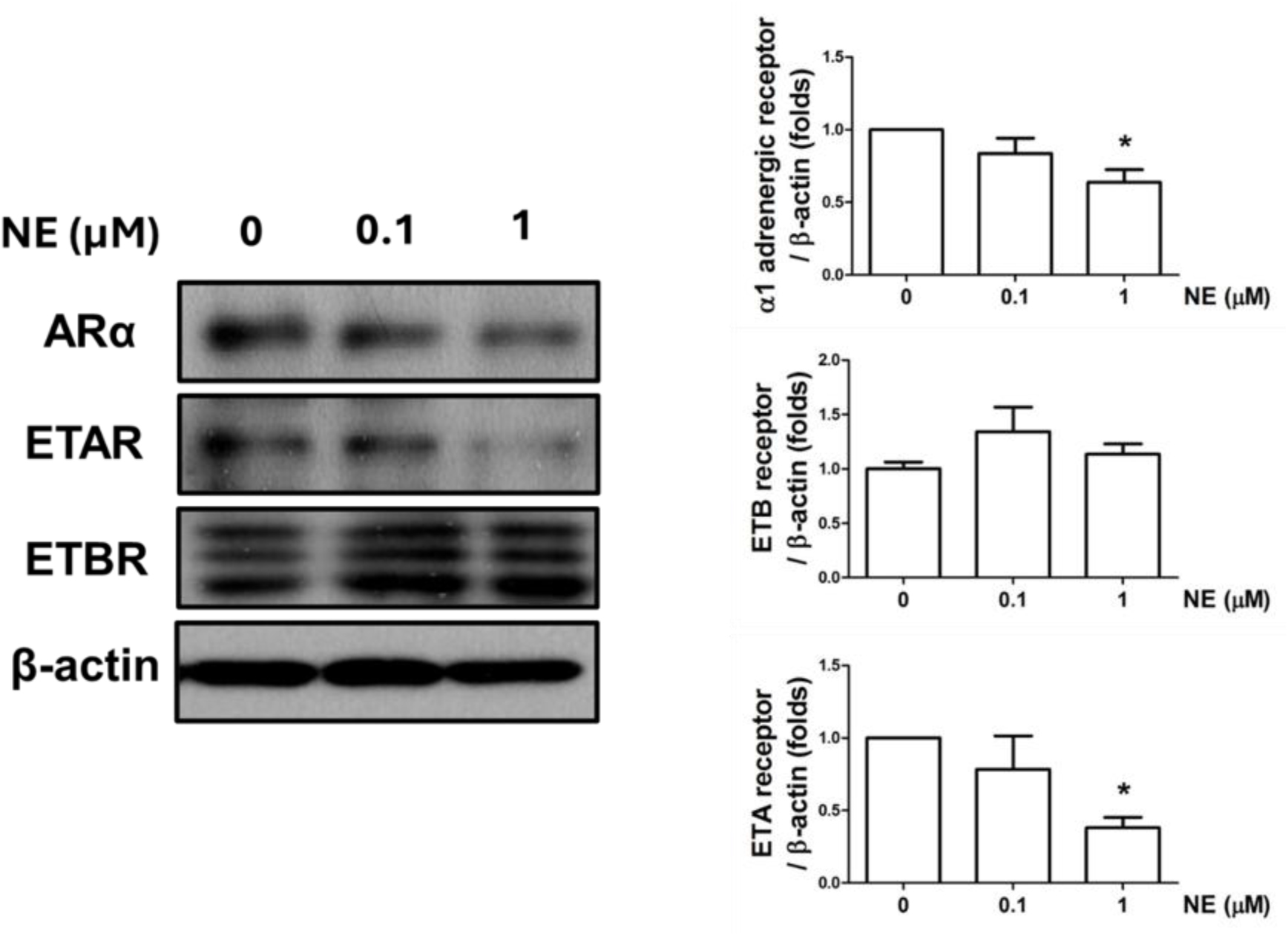
Noradrenalin (NE) decreased the amount of α1 adrenergic receptor (AR α)and Endothelin A receptor (ETA). After 0.1 and 1 μM NE treatment for 6hr, the protein expression of AR α, ETA and ETB was evaluated by western blots. The protein expressions were quantified using densitometric analysis. Results are expressed as means±SEM for at least three independent experiments. *P < 0.05 compared to control group.

**Figure 4.**
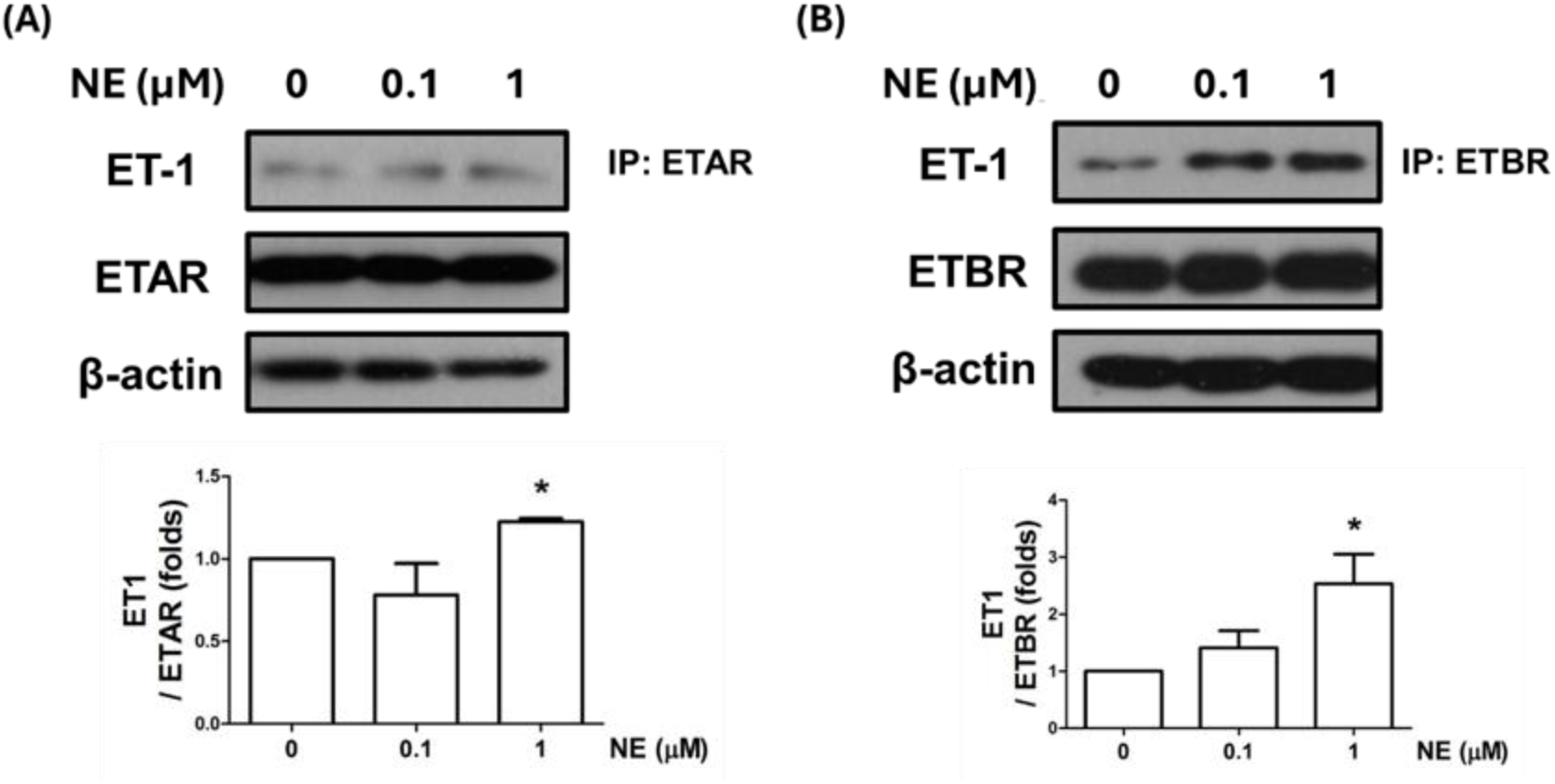
Noradrenalin (NE) induced ET1 to bind the ETA or ETB receptors. After NE treatment for 6hr, equal amounts of protein extracts were subjected to immunoprecipitation (IP) by ETA (A) or ETB (B)receptor antibody. The immunoprecipitates were analyzed by immunoblotting using ET1 antibody (A and B). The protein expressions were quantified using densitometric analysis. Results are expressed as means±SEM for at least three independent experiments. *P < 0.05 compared to control group.

There are several pathways are known for regulating PASMC apoptosis or proliferation. According to previous study, P38 pathway participated in NE-induced rat PASMC proliferation^18^. In this study, we wanted to further clarify whether NE could influence other apoptosis or proliferation-related pathways. After treating 0.1 and 1μM NE for 6hr, the phosphorylation of ERK decreased significantly in a dose-dependent manner (Figure 5). However, the protein level of ERK was not affected by NE treatment. NE treatment caused opposite effects on JNK/c-jun pathway. 1μM NE significantly increased the phosphorylation of JNK and the protein expression of c-jun (Figure 5). The inhibition of ERK pathway and the activation of JNK/c-jun pathway may play an important role in NE-induced PASMC proliferation. Combined previous study and our data could establish a potential action mode of NE on PASMC proliferation induction.

**Figure 5.**
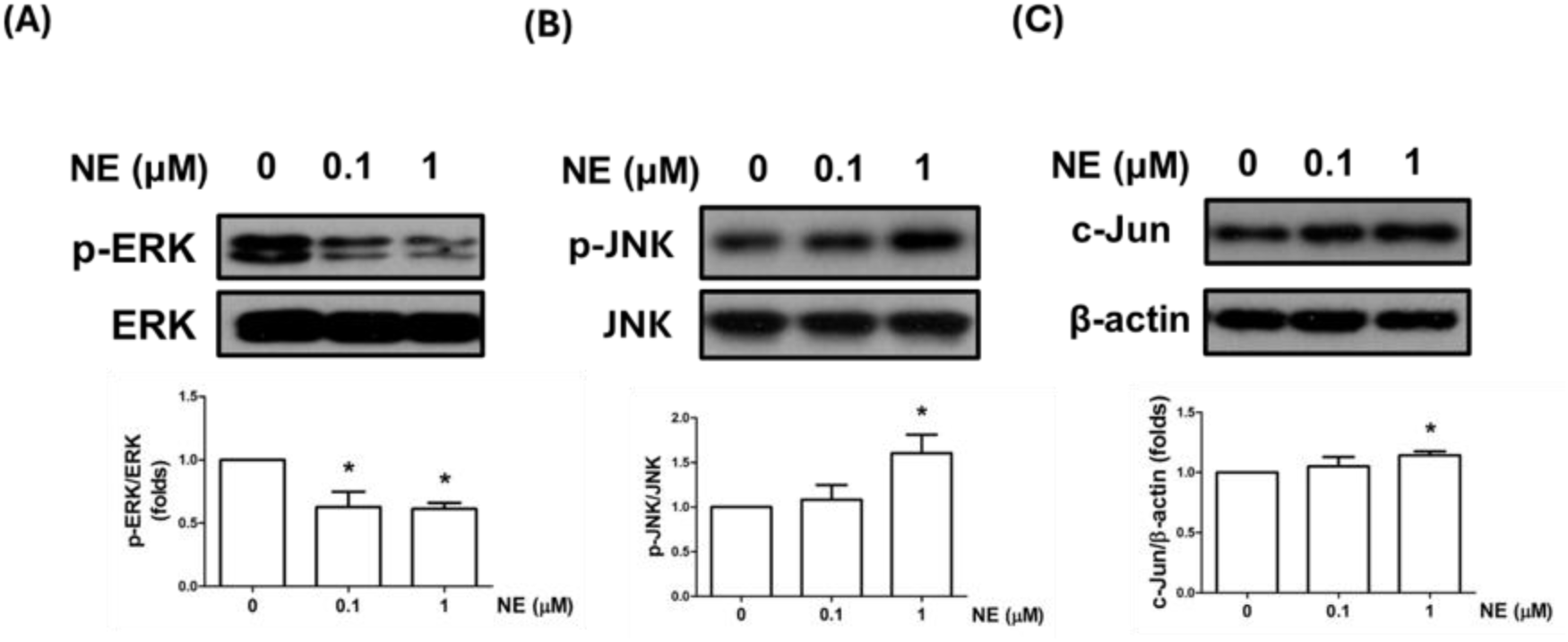
Noradrenalin (NE) triggered the protein expressions of JNK/c-jun pathway. After NE treatment for 6hr, the protein phosphorylation or expressions of ERK (A), JNK (B) and c-Jun (C) were determined by Western blotting. The protein expressions were quantified using densitometric analysis. Results are expressed as means±SEM for at least three independent experiments. *P < 0.05 compared to control group.

## Discussion

Until now, PAH is still a server disease that happened both on adult and infant without an effective therapeutic strategy. PASMC proliferation is a critical pathological event in PAH development. In this study, we investigated that the NE could induced PASMC proliferation and migration. Furthermore, we first find that ET-1 signaling pathway plays an important role in NE-induced PASMC proliferation. From the data on Immunoprecipitation, we also found NE could increase NE-1 secretion, further enhancing ET-1 bind to its receptors. Furthermore, we found that JNK/c-jun pathway was activated after NE treatment which may also participate in NE-induced PASMC proliferation. By combining the results from previous studies, we could better understand the role and mechanism of NE in PASMC proliferation.

There are several studies that investigated the interaction between NE and ET-1 in cardiovascular systems from different perspectives. Most of these studies were focused on ET-1-regulated NE function. ET-1 was considered to enhance NE-induced cardiovascular constriction under normal physical condition and hypertension models^28–30^. ET-1 could regulate NE reuptake or exocytosis through ETAR and ETBR. Kita et al. found that ET-1 enhances NE-induced vascular constriction through ETBR. Another study indicated the function of ET-1 on NE-induced heart constriction is biphasic. ET-1/ETAR pathway could maintain the effect of NE by inhibiting NE uptake, but ET-1/ETBR pathway decrease the level of NE by inhibiting NE release. The final consequence caused by ET-1 depends on the balance between different pathways. There are few studies focused on the effects of NE on ET-1-indcued vascular remodeling. Dao et al. indicated endothelin is an important factor on NE-induced rat vascular hyperplasia^31^, however, the related mechanisms are still unknown. In this study, we first found that NE could induce ET-1 expression and secretion which is one of the important pathways to enhance PASMC proliferation. Our results showed the interaction between NE and ET-1 during PAH development may be bidirectional and complex.

In our results, we find that 0.1-1 μM NE treatment induced the stronger effect of cell proliferation than 5-10 μM NE treatment. This result is consistent with the increase of ET-1. 0.1-1 μM NE treatment induced ET-1 expression significantly, however, the expression of ET-1 in 5-10 μM NE treatment group is lower than control. Furthermore, we observed higher concentrations of NE still could partly enhance PASMC proliferation even though the amount of ET-1 did not increase under the higher concentrations of NE. We believed that the effect of NE on PASMC proliferation should be the balance between different pathways. Previous studies showed that NE could induce PASMC proliferation through p38 and ERK pathways^18, 32^. Besides the ET-1 pathway, platelet Derived Growth Factor BB (PDGF-BB) pathway is another critical pathway to regulate PASMC proliferation during PAH development. Bobik et al also found that NE could activate PDGF-BB-induced DNA synthesis in vascular smooth muscle cells^33^. Furthermore, we believe that there may be a negative feedback regulation between PDGF and ET-1 pathways, because we found that PDGF-BB treatment could inhibit ET-1 mRNA and protein expression in PASMC (data not shown). Even though ET-1 was not increased in 5-10 μM NE treatment group, NE might promote PASMC proliferation through other mechanisms, like PDGF-BB pathway. On the other hand, NE does not always play the role of promoting cell proliferation. Hu et al. found that NE could inhibit TGF-β-induced vascular smooth cell proliferation during aorta remodeling^34^. These studies showed that NE-regulated cell proliferation by multiple-pathways. From our results, we believed ET-1 pathway play a critical role on NE-induced PAMSC proliferation during PAH development. However, depending on different physiological or pathological microenvironments, the effects of NE on PASMC may also change due to the combined results from the different mechanisms.

In this study, we found that NE could induce ET-1 expression and activate JNK/c-jun pathway. There are few study that discussed the cross talk between ET-1 and JNK/c-jun pathway in PASMC. Kyaw et al found that cleaved ET-1 induced JNK activation in rat aortic smooth muscle cells. They also indicated that JNK/AP-1 pathway may participate in ET-1-induced smooth muscle cells proliferation^35^. Furthermore, in astrocyte, ET-1 induced cell proliferation through JNK/c-Jun pathway. These two studies showed that JNK/c-Jun may participate in NE-induced PASMC proliferation through ET-1 pathway. Besides ET-1, JNK/c-jun pathway could involve in PASMC proliferation through other signaling pathways during PAH development, such as PDGF-BB pathway. Zhao et al indicated the PASMC proliferation induced by PDGF-BB is JNK-dependent, but not p38 or ERK^36^. MiRNA-4632 could inhibit PDGF-BB-induced human PASMC proliferation by targeting c-jun directly^37^. Furthermore, several studies showed that JNK signaling is the major mechanism of different compounds that attenuated PDGF-BB-induced PASMC proliferation^38, 39^. In this study, we did not clarify whether NE could interfere with the PDGF-BB-induced PASMC proliferation. However, a study in 1990 could provide the supported information. Bobik et al proved that NE could enhance the binding efficiency of PDGF-BB and its receptor^33^. Combined with the information, it is possible that JNK/c-jun pathways participate in NE-induced PASMC proliferation through both ET-1 and PDGF-BB. However, more data is needed to clarify this mechanism.

## Nonstandard Abbreviations and Acronyms

ARα: androgen receptor
ET1: Endothelin 1
ETRA: endothelin-1 receptor A
ETRB: endothelin-1 receptor B
HIF-1α: hypoxia-inducible factor-1α
MAPK: mitogen-activated protein kinases
miRNA: microRNA
MTT: 3-(4,5-dimethyl thiazol-2-yl)-2,5-diphenyltetrazolium bromide
NE: noradrenalin
PAH: pulmonary arterial hypertension
PASMC: pulmonary artery smooth muscle cells
PBST: phosphate-buffered saline with Tween
PDGF-BB: platelet Derived Growth Factor BB
SGB: sympathetic ganglion blockage
SMC: smooth muscle cell

## Acknowledgments

The authors thank National Taiwan University Hospital and National Taiwan University Children’s Hospital for technical and funding support.

## Sources of Funding

This study was supported by grants from NTUH. 113-S0179 and NTUH.112-S0189.

## Disclosures

None

## Notes

### Competing Interest Statement

The authors have declared no competing interest.

## References

1. Cassin S, Dawes GS, Mott JC, Ross BB and Strang LB. THE VASCULAR RESISTANCE OF THE FOETAL AND NEWLY VENTILATED LUNG OF THE LAMB. The Journal of physiology. 1964;171:61–79.

2. Weir EK and Archer SL. The mechanism of acute hypoxic pulmonary vasoconstriction: the tale of two channels. FASEB journal : official publication of the Federation of American Societies for Experimental Biology. 1995;9:183–9.

3. Resnik ER, Herron JM, Lyu SC and Cornfield DN. Developmental regulation of hypoxia-inducible factor 1 and prolyl-hydroxylases in pulmonary vascular smooth muscle cells. Proceedings of the National Academy of Sciences of the United States of America. 2007;104:18789–94.

4. Ball MK, Waypa GB, Mungai PT, Nielsen JM, Czech L, Dudley VJ, Beussink L, Dettman RW, Berkelhamer SK, Steinhorn RH, Shah SJ and Schumacker PT. Regulation of hypoxia-induced pulmonary hypertension by vascular smooth muscle hypoxia-inducible factor-1α. American journal of respiratory and critical care medicine. 2014;189:314–24.

5. Kapitsinou PP, Rajendran G, Astleford L, Michael M, Schonfeld MP, Fields T, Shay S, French JL, West J and Haase VH. The Endothelial Prolyl-4-Hydroxylase Domain 2/Hypoxia-Inducible Factor 2 Axis Regulates Pulmonary Artery Pressure in Mice. Molecular and cellular biology. 2016;36:1584–94.

6. Dai Z, Li M, Wharton J, Zhu MM and Zhao YY. Prolyl-4 Hydroxylase 2 (PHD2) Deficiency in Endothelial Cells and Hematopoietic Cells Induces Obliterative Vascular Remodeling and Severe Pulmonary Arterial Hypertension in Mice and Humans Through Hypoxia-Inducible Factor-2α. Circulation. 2016;133:2447–58.

7. Kim YM, Barnes EA, Alvira CM, Ying L, Reddy S and Cornfield DN. Hypoxia-inducible factor-1α in pulmonary artery smooth muscle cells lowers vascular tone by decreasing myosin light chain phosphorylation. Circulation research. 2013;112:1230–3.

8. Maruyama H, Sakai S and Ieda M. Endothelin-1 alters BMP signaling to promote proliferation of pulmonary artery smooth muscle cells. Canadian journal of physiology and pharmacology. 2022;100:1018–1027.

9. Kim FY, Barnes EA, Ying L, Chen C, Lee L, Alvira CM and Cornfield DN. Pulmonary artery smooth muscle cell endothelin-1 expression modulates the pulmonary vascular response to chronic hypoxia. American journal of physiology Lung cellular and molecular physiology. 2015;308:L368–77.

10. Jankov RP, Kantores C, Belcastro R, Yi M and Tanswell AK. Endothelin-1 inhibits apoptosis of pulmonary arterial smooth muscle in the neonatal rat. Pediatric research. 2006;60:245–51.

11. Nadeau V, Potus F, Boucherat O, Paradis R, Tremblay E, Iglarz M, Paulin R, Bonnet S and Provencher S. Dual ET(A)/ET(B) blockade with macitentan improves both vascular remodeling and angiogenesis in pulmonary arterial hypertension. Pulmonary circulation. 2018;8:2045893217741429.

12. Li M, Liu Y, Jin F, Sun X, Li Z, Liu Y, Fang P, Shi H and Jiang X. Endothelin-1 induces hypoxia inducible factor 1α expression in pulmonary artery smooth muscle cells. FEBS letters. 2012;586:3888–93.

13. Pisarcik S, Maylor J, Lu W, Yun X, Undem C, Sylvester JT, Semenza GL and Shimoda LA. Activation of hypoxia-inducible factor-1 in pulmonary arterial smooth muscle cells by endothelin-1. American journal of physiology Lung cellular and molecular physiology. 2013;304:L549–61.

14. Na S, Kim OS, Ryoo S, Kweon TD, Choi YS, Shim HS and Oh YJ. Cervical ganglion block attenuates the progression of pulmonary hypertension via nitric oxide and arginase pathways. Hypertension (Dallas, Tex : 1979). 2014;63:309–15.

15. Salvi SS. Alpha1-adrenergic hypothesis for pulmonary hypertension. Chest. 1999;115:1708–19.

16. Zhang H and Faber JE. Trophic effect of norepinephrine on arterial intima-media and adventitia is augmented by injury and mediated by different alpha1-adrenoceptor subtypes. Circulation research. 2001;89:815–22.

17. Ye C, Zheng F, Xu T, Wu N, Tong Y, Xiong XQ, Zhou YB, Wang JJ, Chen Q, Li YH, Zhu GQ and Han Y. Norepinephrine acting on adventitial fibroblasts stimulates vascular smooth muscle cell proliferation via promoting small extracellular vesicle release. Theranostics. 2022;12:4718–4733.

18. Xiao X, Zhang Y, Tian S, Wang X, Zhang Q, Zhang L, Yu X, Ma C, Zheng X, Li Y, Zhang J and Qu L. Alpha1B-adreneroceptor is involved in norepinephrine-induced pulmonary artery smooth muscle cell proliferation via p38 signaling. European journal of pharmacology. 2022;931:175159.

19. Wang Y, Ruan Y and Wu S. ET-1 regulates the human umbilical vein endothelial cell cycle by adjusting the ERβ/FOXN1 signaling pathway. Annals of translational medicine. 2020;8:1499.

20. Spinella F, Rosanò L, Di Castro V, Decandia S, Nicotra MR, Natali PG and Bagnato A. Endothelin-1 and endothelin-3 promote invasive behavior via hypoxia-inducible factor-1alpha in human melanoma cells. Cancer research. 2007;67:1725–34.

21. Wu MH, Chen LM, Hsu HH, Lin JA, Lin YM, Tsai FJ, Tsai CH, Huang CY and Tang CH. Endothelin-1 enhances cell migration through COX-2 up-regulation in human chondrosarcoma. Biochimica et biophysica acta. 2013;1830:3355–64.

22. Cheng X, Yeung PKK, Zhong K, Zilundu PLM, Zhou L and Chung SK. Astrocytic endothelin-1 overexpression promotes neural progenitor cells proliferation and differentiation into astrocytes via the Jak2/Stat3 pathway after stroke. Journal of neuroinflammation. 2019;16:227.

23. Yen TA, Huang HC, Wu ET, Chou HW, Chou HC, Chen CY, Huang SC, Chen YS, Lu F, Wu MH, Tsao PN and Wang CC. Microrna-486-5P Regulates Human Pulmonary Artery Smooth Muscle Cell Migration via Endothelin-1. International journal of molecular sciences. 2022;23.

24. Clozel M and Salloukh H. Role of endothelin in fibrosis and anti-fibrotic potential of bosentan. Annals of medicine. 2005;37:2–12.

25. Weng CM, Yu CC, Kuo ML, Chen BC and Lin CH. Endothelin-1 induces connective tissue growth factor expression in human lung fibroblasts by ETAR-dependent JNK/AP-1 pathway. Biochemical pharmacology. 2014;88:402–11.

26. Yamboliev IA, Hruby A and Gerthoffer WT. Endothelin-1 activates MAP kinases and c-Jun in pulmonary artery smooth muscle. Pulmonary pharmacology & therapeutics. 1998;11:205–8.

27. Kirkby NS, Hadoke PW, Bagnall AJ and Webb DJ. The endothelin system as a therapeutic target in cardiovascular disease: great expectations or bleak house? British journal of pharmacology. 2008;153:1105–19.

28. Kita S, Taguchi Y and Matsumura Y. Endothelin-1 enhances pressor responses to norepinephrine: involvement of endothelin-B receptor. Journal of cardiovascular pharmacology. 1998;31 Suppl 1:S119–21.

29. Henrion D and Laher I. Potentiation of norepinephrine-induced contractions by endothelin-1 in the rabbit aorta. Hypertension (Dallas, Tex : 1979). 1993;22:78–83.

30. Kita S, Taguchi Y, Chatani S and Matsumura Y. Effects of endothelin-1 on norepinephrine-induced vasoconstriction in deoxycorticosterone acetate-salt hypertensive rats. European journal of pharmacology. 1998;344:53–7.

31. Dao HH, Lemay J, de Champlain J, deBlois D and Moreau P. Norepinephrine-induced aortic hyperplasia and extracellular matrix deposition are endothelin-dependent. Journal of hypertension. 2001;19:1965–73.

32. Liu R, Zhang Q, Luo Q, Qiao H, Wang P, Yu J, Cao Y, Lu B and Qu L. Norepinephrine stimulation of alpha1D-adrenoceptor promotes proliferation of pulmonary artery smooth muscle cells via ERK-1/2 signaling. The international journal of biochemistry & cell biology. 2017;88:100–112.

33. Bobik A, Grinpukel S, Little PJ, Grooms A and Jackman G. Angiotensin II and noradrenaline increase PDGF-BB receptors and potentiate PDGF-BB stimulated DNA synthesis in vascular smooth muscle. Biochemical and biophysical research communications. 1990;166:580–8.

34. Hu Z, Li B, Wang Z, Hu X, Zhang M, Chen R, Wu Q and Jia F. The sympathetic transmitter norepinephrine inhibits VSMC proliferation induced by TGFβ by suppressing the expression of the TGFβ receptor ALK5 in aorta remodeling. Molecular medicine reports. 2020;22:387–397.

35. Kyaw M, Yoshizumi M, Tsuchiya K, Kirima K, Suzaki Y, Abe S, Hasegawa T and Tamaki T. Antioxidants inhibit endothelin-1 (1-31)-induced proliferation of vascular smooth muscle cells via the inhibition of mitogen-activated protein (MAP) kinase and activator protein-1 (AP-1). Biochemical pharmacology. 2002;64:1521–31.

36. Zhao Y, Lv W, Piao H, Chu X and Wang H. Role of platelet-derived growth factor-BB (PDGF-BB) in human pulmonary artery smooth muscle cell proliferation. Journal of receptor and signal transduction research. 2014;34:254–60.

37. Qian Z, Li Y, Chen J, Li X and Gou D. miR-4632 mediates PDGF-BB-induced proliferation and antiapoptosis of human pulmonary artery smooth muscle cells via targeting cJUN. American journal of physiology Cell physiology. 2017;313:C380–c391.

38. Chang Y, Uen YH, Chen CC, Lin SC, Tseng SY, Wang YH, Sheu JR and Hsieh CY. Platonin inhibited PDGF-BB-induced proliferation of rat vascular smooth muscle cells via JNK1/2-dependent signaling. Acta pharmacologica Sinica. 2011;32:1337–44.

39. Yang PS, Wang MJ, Jayakumar T, Chou DS, Ko CY, Hsu MJ and Hsieh CY. Antiproliferative Activity of Hinokitiol, a Tropolone Derivative, Is Mediated via the Inductions of p-JNK and p-PLCγ1 Signaling in PDGF-BB-Stimulated Vascular Smooth Muscle Cells. Molecules (Basel, Switzerland). 2015;20:8198–212.

